# Sexual behavior of the desert locust during intra-and inter-phase interactions

**DOI:** 10.1101/377903

**Authors:** Y. Golov, J. Rillich, M. Douek, A.R. Harari, A Ayali

## Abstract

Mating and reproduction behaviors and strategies are fundamental aspects of an organism’s evolutionary and ecological success. In locusts, intra- as well as inter-phase reproductive interactions among gregarious and solitarious locust populations have a major impact on the locust population dynamics. However, practically all previous work on locust sexual behavior has been limited to the gregarious phase. Here we provide a first detailed description of pre-copulatory behavior of solitarious desert locusts. We compare our findings with those of previous reports of pre-copulatory behavior of gregarious locusts, focusing on the behavioral elements that serve in inter-sex signaling and communication. We also studied inter-phase (mixed pairs) reproductive interactions. Solitarious males were found to invest more in pre-copulatory courtship and signaling compared to their gregarious counterparts; and the solitarious females played a comparatively more dominant role in the inter-sex communication. The solitarious females were also less prone to demonstrate the typical rejection-related behavioral patterns displayed by the gregarious females. As a consequence of the particular characteristic behavior of each phase, the most successful among intra- and inter-phase pairs were gregarious males with solitary females. Least successful were solitary males encountered with gregarious females, indicating a strong asymmetry in inter-phase reproductive interactions. We discuss these results in the context of non-random or assortative mating in locust mixed or sympatric solitarious-gregarious populations.

## Introduction

The desert locust (*Schistocerca gregaria*) presents a remarkable example of environmentally-induced behavioral plasticity: changes in population density dramatically alter this locust’s behavior. The major behavioral characteristic of locusts in the gregarious phase is that of their strong attraction to conspecifics, which translates to active aggregation behavior (Ellis, 1959; Ellis, 1963; Uvarov, 1966). Gregarious locusts are also generally more active, featuring a strong propensity to march in huge bands of hoppers and form high-density swarms of reproductive adults (recent reviews in Ariel & Ayali, 2015; Cullen *et al.*, 2017). In contrast, solitary-reared locusts (or locusts from solitarious field populations) actively avoid contact with other locusts (Roessingh *et al.*, 1993; Geva *et al.*, 2010), are more sedentary and cryptic in behavior, do not march, and fly less. Phase transformation is a positive-feedback process. The importance of the change in behavior is that it precedes and facilitates all other phase-changes, from biochemistry and physiology to coloration and morphology (Ariel & Ayali, 2015; Cullen *et al.*, 2017).

Similar to other species demonstrating population cycles (or fluctuations), key factors that affect the dynamics of locust populations are those of mating and reproduction strategies and behaviors. Hence, reproductive success is closely tied to the risks of locust outbreaks (Farrow, 1979; Topaz *et al.*, 2012). Acquiring an understanding of these dynamics calls for an initial in-depth knowledge of the mating and reproductive behavior of locusts. Locust courtship, in general, is considered simple and primitive (Loher, 1959; Uvarov, 1966, 1977; Strong & Amerasinghe, 1977). Males of *S. gregaria* were reported to be more dominant during sexual-interactions, while females were reported to show no clear courtship behavioral patterns (Norris, 1954; Pener, 1965, 1967; Strong & Amerasinghe, 1977; Amerasinghe, 1978; Inayatullah *et al.*, 1994; Golov *et al.*, 2018). A major feature of the female locust sexual behavior was reported to be rejection of the males: including jumping (before and during mounting), kicking, and movements of the abdomen to prevent copulation (Loher, 1959; Strong & Amerasinghe, 1977; Uvarov, 1977; Golov *et al.*, 2018). We recently provided a first comprehensive description of the pre-copulatory behavior of the gregarious desert locust (Golov *et al.*, 2018), presenting a detailed quantification of the sexual behavior of both sexes, with special emphasis on female choice.

The above-noted study by Golov et al. (2018), however, as well as most of the above-mentioned earlier work, was limited to locusts in the gregarious phase (but see Inayatullah *et al.*, 1994). In the gregarious phase, the desert locust displays clear sexual dimorphism, in which fully mature males are bright yellow and females beige-brown to yellowish (Norris, 1954; Pener, 1965; Pener & Simpson, 2009). In accordance with this, it has been assumed that visual cues are a primary means of sexual recognition and pre-copulatory communication in the gregarious locust (Whitman, 1990; Obeng-Ofori *et al.*, 1993; Inayatullah *et al.*, 1994; Ely *et al.*, 2006). Sexual dimorphism is much less pronounced in the solitarious phase of the desert locust. Very limited knowledge is available regarding sexual recognition and communication, as well as courtship and mating behavior of locusts in the solitarious phase. For example, Tanaka et al. (2003) reported that in isolated-reared adult migratory locusts (*Locusta migratoria*) the pre-copulatory mounting tends to be shorter than that of crowd-reared ones, while the opposite is true for the length of copulation. In the desert locust, as in all true locust species, the fundamental differences in life history between the two phases are expected to result in major differences in courtship and pre-copulatory behavior. Density-dependent phase differences can be predicted in all stages and aspects of the behavior, from mate finding and mate recognition, through pre-mounting courtship, to mate choice and sexual conflict.

A practically unexplored question of interest, as well as of importance for our understanding of locust population dynamics, is that of the existence and nature of inter-phase reproductive interactions. Locust populations of mixed phase or, alternatively, sympatric populations of gregarious and solitarious locusts, can be found during periods of transition, i.e. in recession areas, during periods of increasing populations, or in migration areas during the breakdown of a swarm (e.g. Tanaka *et al.*, 2010). As noted, density-dependent variations are expected in locust reproductive behavior. Alternative reproductive strategies or tactics are accepted to be aimed at increasing the players’ relative reproductive success. However, the interactions of some reproductive phenotypes may restrict gene flow within the species (e.g. assortative mating; Andersson & Iwasa, 1996; Nosil *et al.*, 2005; Rundle & Nosil, 2005; Ramirez-Romero *et al.*, 2017). In the case of phase-dependent reproductive phenotypes in locusts, this may translate to limited inter-phase pre-reproductive interactions and an overall preference to mate within the phase.

In the current work we set out to present a comparative, detailed description of pre-copulatory behavior of male and female desert locusts in the solitarious phase. We further examined how the pronounced phase-dependent differences affect inter-phase encounters and the success of inter-phase copulation. Our findings have important consequences for understanding locust population dynamics, and specifically the dynamics of phase change and the development of a locust outbreak.

## Material & Methods

### Animals

Desert locusts, *Schistocerca gregaria* (Forskål 1775), from our colony at Tel Aviv University were reared for many consecutive generations under crowded conditions (i.e. approaching the gregarious phase), 100-160 individuals in 60L aluminum cages. In order to obtain locusts in the solitarious phase, hatchlings from eggs laid by the gregarious females were collected within 2 h of hatching and reared in isolation in 1L cages until adulthood (Geva *et al.*, 2010). Extra care was taken to maintain similar rearing conditions for both gregarious and solitarious locusts, with the exception of density. Cages were located in two dedicated room (separating between gregarious and solitarious colonies) under a constant temperature (29-31°C) and light cycle of 12L:12D. Supplementary radiant heat was supplied during day-time by incandescent 25 W electric bulbs (full visible spectrum, yellow and red dominant), resulting in a day temperature of ca. 37°C. Locusts were provided daily with fresh wheat and dry oats, and plastic cups filled with moist sand for oviposition.

Adult locusts used in all experiments were virgin males and females. Gregarious virgin adults were obtained by marking newly-emerged adults with non-poisonous acrylic paint within 24 hours following ecdysis. Males and females were separated into single- sex “cohort cages” every 3 days. Thus, in each cohort cage the maximum age range of the individual locusts was less than 72 hours. The cages were maintained under the same rearing conditions as above. Based on our preliminary work and previous reports (Injeyan & Tobe, 1981; Mahamat *et al.*, 1993; Wybrandt & Andersen, 2001; Ely *et al.*, 2006; Nishide & Tanaka, 2012), gregarious locusts in our experiments were sexually mature at 12-14-days-old in males (when the yellowish coloration had reached stage V; see Norris, 1954; Loher, 1961), and at 18-20 days-old in females. The maturation period is longer in solitarious locusts than in their gregarious counterparts (Norris, 1954; Norris, 1962; Pener, 1976; Amerasinghe, 1978)). Hence, the age of solitarious experimental animals of both sexes was 30-32 days.

### Experimental design

Experiments were carried out as described by Golov et al. (2018). Briefly, a male and a female locust, in the solitary phase, were introduced into the two sides of an observation cell (14×13×24 cm, divided in two by an opaque plastic partition). Five minutes after introducing the locusts into the cell the experiment was initiated by carefully removing the partition between the compartments, and replacing an initial red light illumination with two regular 25 W light bulbs. Experiments lasted up to 3 hours, or until copulation had occurred, if earlier, and were recorded by a SONY HDR-PJ820E video camera.

Mixed inter-phase experiments examining the interactions between the phases were conducted in a similar manner as the above, with either a solitary female and a gregarious male or a gregarious female and a solitary male introduced into the cage.

### Data analyses

The recorded videos of the behavior of each pair were reviewed and analyzed using BORIS software (Friard & Gamba, 2016). Data analyses mostly followed the detailed description provided by Golov et al. (2018). In brief, behavioral elements were identified in order to describe the locusts’ pre-copulatory behavior. These included both repetitive (prolonged, e.g. the vibration of the hind leg femur) and discrete (momentary, e.g. jumping) behaviors. The two behavioral types were counted, with a ‘count’ relating to the duration of a behavior from initiation until termination. Behavioral measurements were taken only if the male and female were at a distance of less than 10 cm (depicted as an ‘encounter’). For both pre-mounting and mounting behavioral elements, the following parameters were measured and compared for both sexes: (1) In order to obtain the pattern or chronological sequence of the behavioral repertoire, the relative latency to initiation of each behavior was noted (relative to the total time of the relevant stage, either pre-mounting or mounting). (2) The probability of a specific behavior occurring (per experiment, PO=1 if the behavior occurred at least once, and 0 otherwise). (3) The frequency of occurrences of a specific behavioral element.

Kinematic diagrams were constructed for all the transitions between pairs of behavioral elements (i.e., preceding-following elements), for males and females, as detailed in (Golov *et al.*, 2018). An average transition probability (TP) was calculated based on all possible transitions between each pair of elements. For values of TP between two elements above 10%, a difference of 15% or more, in solitary vs. gregarious locusts, was set as a threshold in constructing the comparative transitions diagrams presented.

The statistical data analysis and output were conducted in GraphPad Prism version 6.04 for Windows, JMP®, Version 12.0.1 SAS Institute, and Canvas draw 2.0 (Deneba Systems, Miami, FL). Comparison were conducted using two-sided Fisher exact test with Bonferroni adjustments as required (for multiple comparisons).

## Results

We first set out to obtain an in-depth comparative description of the pre-copulatory behavior of male and female locusts in the solitarious phase (N=26 pairs). Figure 1 presents, in a simple and schematic manner, the consecutive stages (S1-7) leading from first encounter to copulation of a solitarious locust pair, divided into the pre-mounting and the mounting stages. In Supplementary Figure 1 we detail the behavioral elements comprising each stage, as well as the probability of each element to occur (PO), separately for males and females, in a similar manner to that used to describe the gregarious locusts’ behavior in Golov et al. (2018) (see Fig.1 therein).

**Figure 1:**
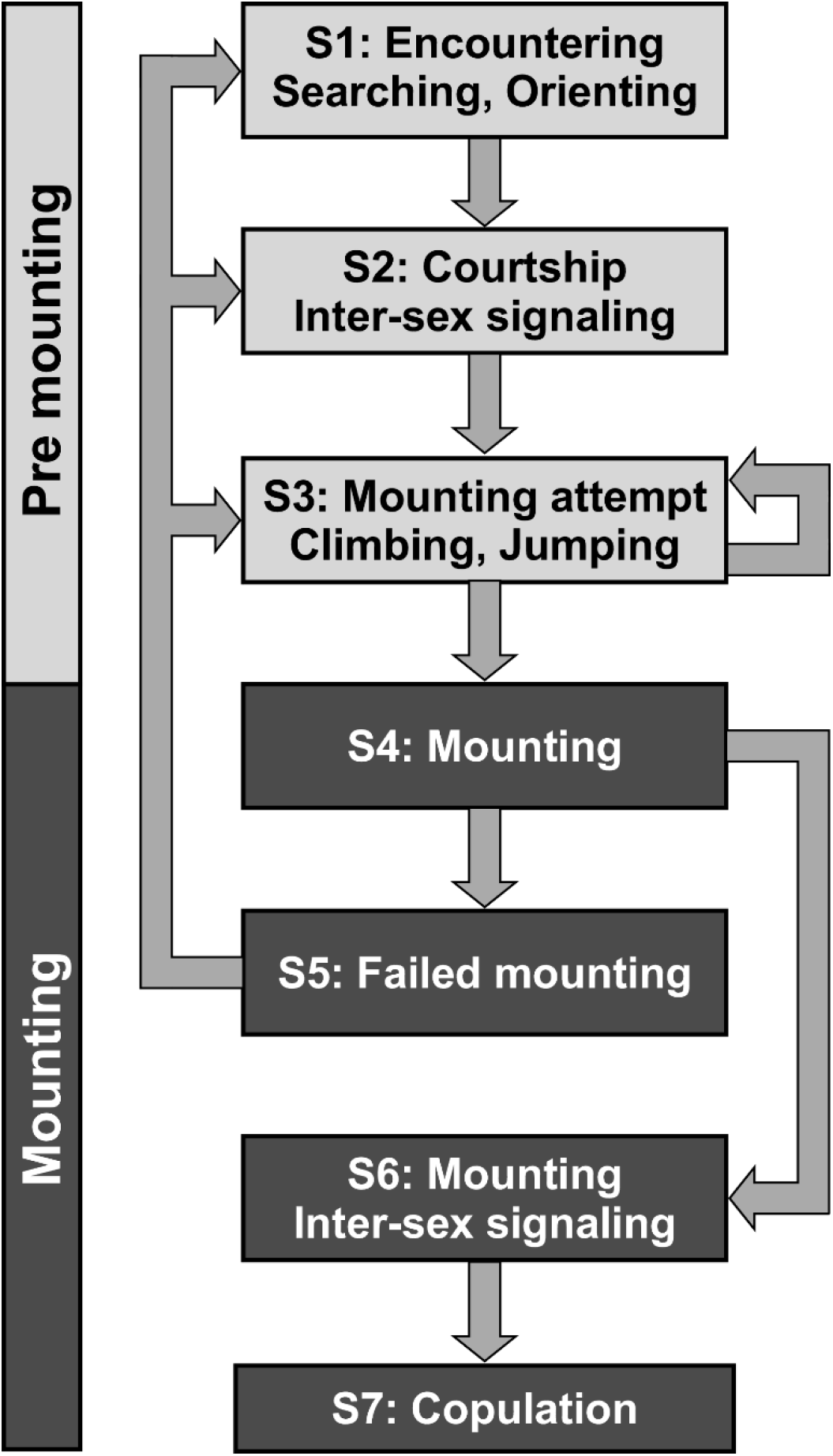
A simplified schematic description of the consecutive stages (S1-7) comprising the solitarious locusts’ pre-copulatory behavioral sequence, from first encounter to copulation. The different stages comprise the premounting and the mounting phases.

In order to facilitate a comparative investigation of phase-related differences in pre-copulatory behavior, we focused our attention on those behavioral elements that we consider fundamental for inter-sex communication (Figs. 2 and 3; raw data for the gregarious locusts are mostly those found in Golov et al., 2018. N=31; Note that the current study included the behavioral data of all pairs tested, unlike Golov et al., 2018 that presented only data from experiments that culminated in successful copulation). Generally, the major phase-related differences were manifested in the pre-mounting stage (cf. Figs 2 and 3 with the different stages noted in Fig 1S). As demonstrated in Figure 2A, solitarious males showed a much higher prevalence of leg vibrations and wing stridulations (see also Figure S2, demonstrating the higher prevalence of transitions to leg and wing signaling in the solitary males). In contrast, a slow repetitive elevation of the hind legs as well as signals associated with the abdomen (abdominal wagging) are behavioral elements much less characteristic of the solitarious males than the gregarious ones. An overall subordinate role of the solitary males in the sexual interactions was manifested by their higher tendency to walk away during pre-mounting (compared to their gregarious counterparts). No phase-related preference was observed for the mode of attempting to mount the female (by way of climbing or jumping; Fig. 2B). Using the wings for signaling (i.e. short and long wing stridulation during pre-mounting) was found to be a characteristic behavior of the solitarious females, almost never demonstrated by gregarious ones (Fig. 3A). A major phase-dependent difference was observed in behavioral elements associated with female rejection, which were much reduced in solitary females (Fig. 3B; Fig S3).

**Figure 2:**
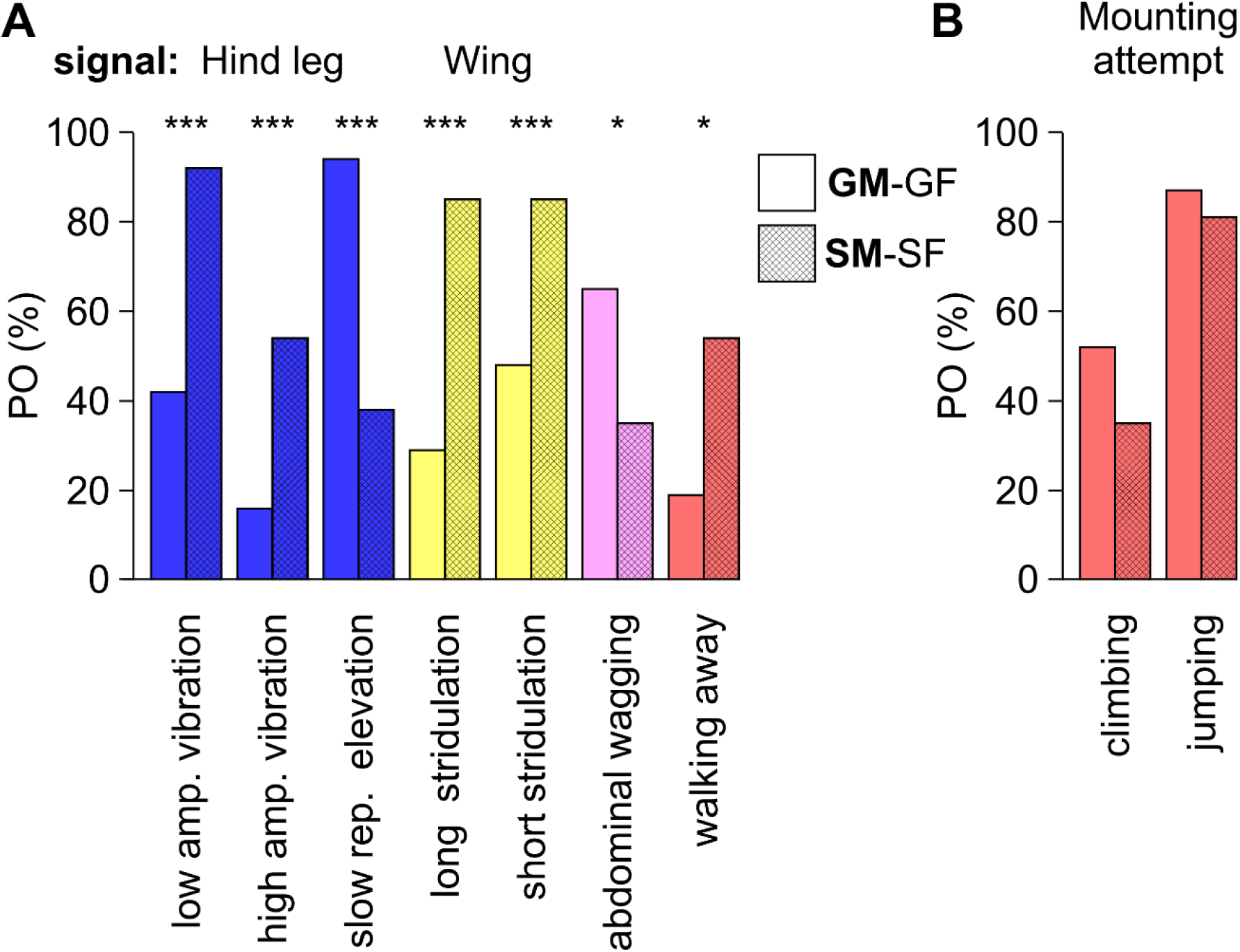
A. The mean probability of occurrence (PO) of several selected male behavioral elements related to sex communication and signaling during pre-mounting. Different colors indicate the body part involved in signaling (indicated on top of bars). B. The mean probability of occurrence (PO) of two types of mounting attempts. Data for solitarious locusts (cross hatched bars, n=26) are shown in comparison to those of gregarious locusts (open bars, n=31; raw data for the gregarious locusts are those found in Golov et al., 2018). *** Statistical significance is indicated: * p<0.05 *** p<0.001.

**Figure 3:**
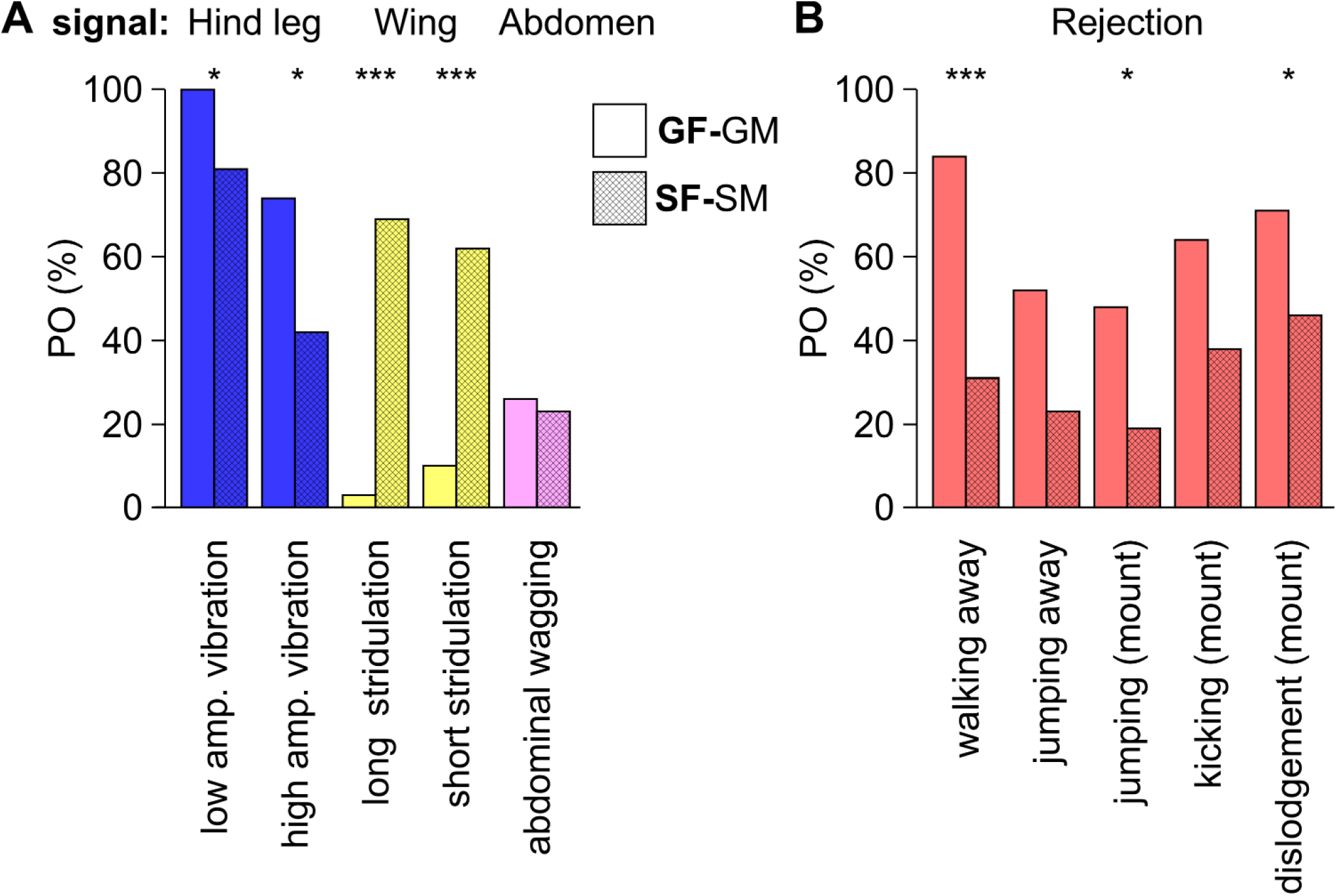
A. Comparison of female pre-mounting behavioral elements presented in a similar way to that in Fig. 2A. B. The mean probability of occurrence (PO) of behavioral elements associated with female pre-copulatory rejection behavior. (mount) denotes after mounting, or during the mounting phase. Colors and bars as detailed in Fig 2. Solitarious, n=26; gregarious, n=31. Raw data for the gregarious locusts are those found in Golov et al., 2018. Statistical significance: * p<0.05 *** p<0.001.

Next, we addressed the nature of the inter-phase reproductive interactions by repeating the experiments presented above, this time with “mixed-phase” pairs. Interestingly, we found that the locusts tended to preserve the dominant features of their sexual behavior, irrespective of the phase-identity of their mate (cf. Figs. 4 & 5 to Figs. 2 & 3, respectively). The solitarious males were still greatly preoccupied with leg vibration and wing stridulation, also when encountering a gregarious female; while the gregarious males still displayed intense slow repetitive elevation of their legs also towards a solitarious female (Fig. 4A). Again, the solitarious females displayed intense wing stridulation, as well as reduced rejection-related behavioral patterns, compared to the gregarious ones (Fig. 5A). Most importantly, the two types of inter-phase pairs differed greatly in the rate of successful copulations (Fig. 6): when paired with a female of the other phase, gregarious males were highly successful; whereas the solitarious males were extremely unsuccessful in inter-phase pairing (9 out of 10 pairs mated successfully compared to only 3 out of 10, respectively; Fig. 6; data for gregarious-gregarious couples were computed from the raw data in Golov et al., 2018). The low rate of successful copulation of the solitary males when paired with gregarious females is further emphasized when compared with the relatively high success rates (more than 80%) demonstrated when interacting with solitarious females.

**Figure 4:**
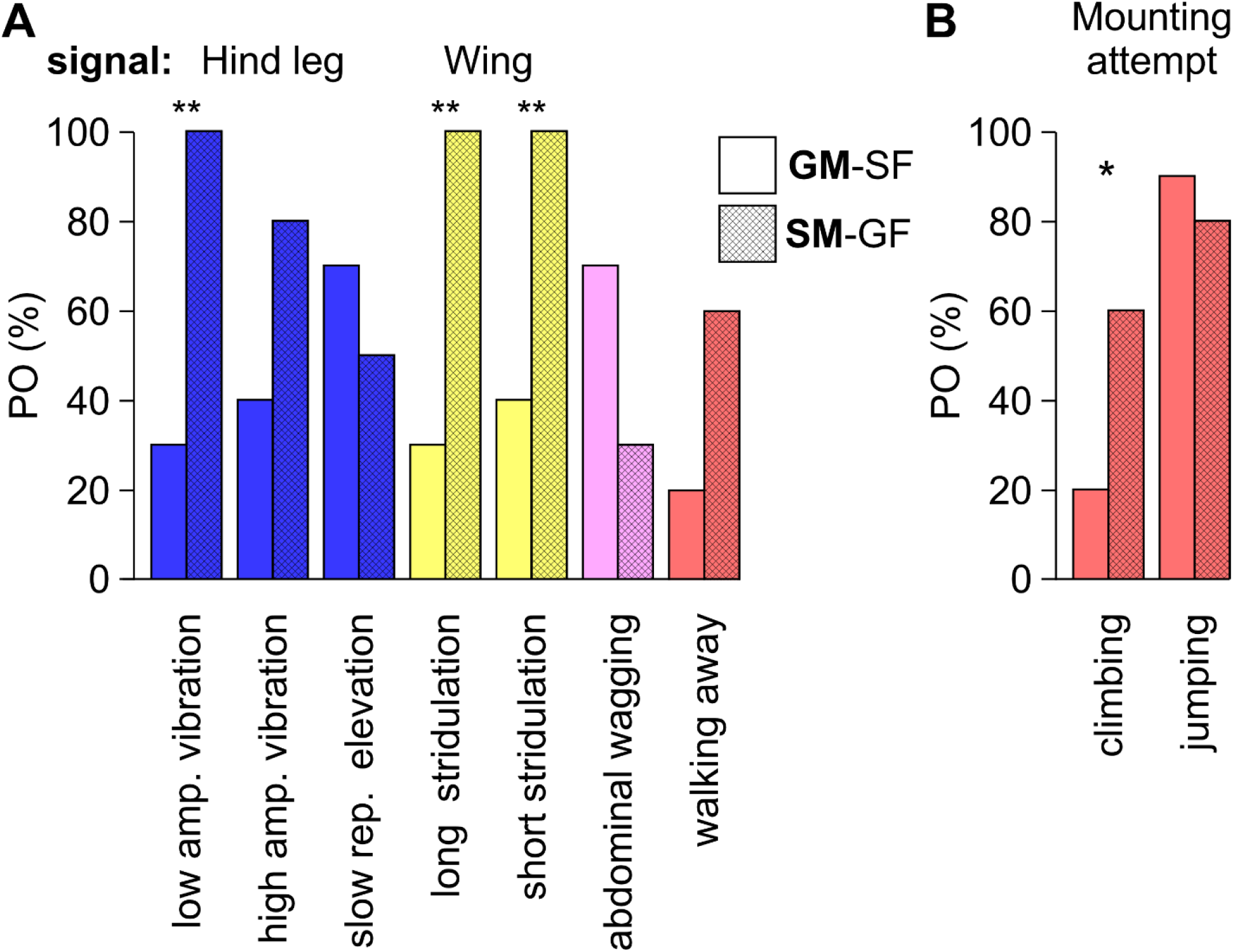
A. Pre-copulatory behavioral elements related to the sexual behavior of males in mixed-phase pairs (GM-SF, Gregarious male and solitarious female; or SM_GF, solitarious male and gregarious female; n=10 for both). Colors and bars as detailed in Fig 2. Raw data for the gregarious locusts are those found in Golov et al., 2018. Statistical significance: * p<0.05 ** p<0.01.

**Figure 5:**
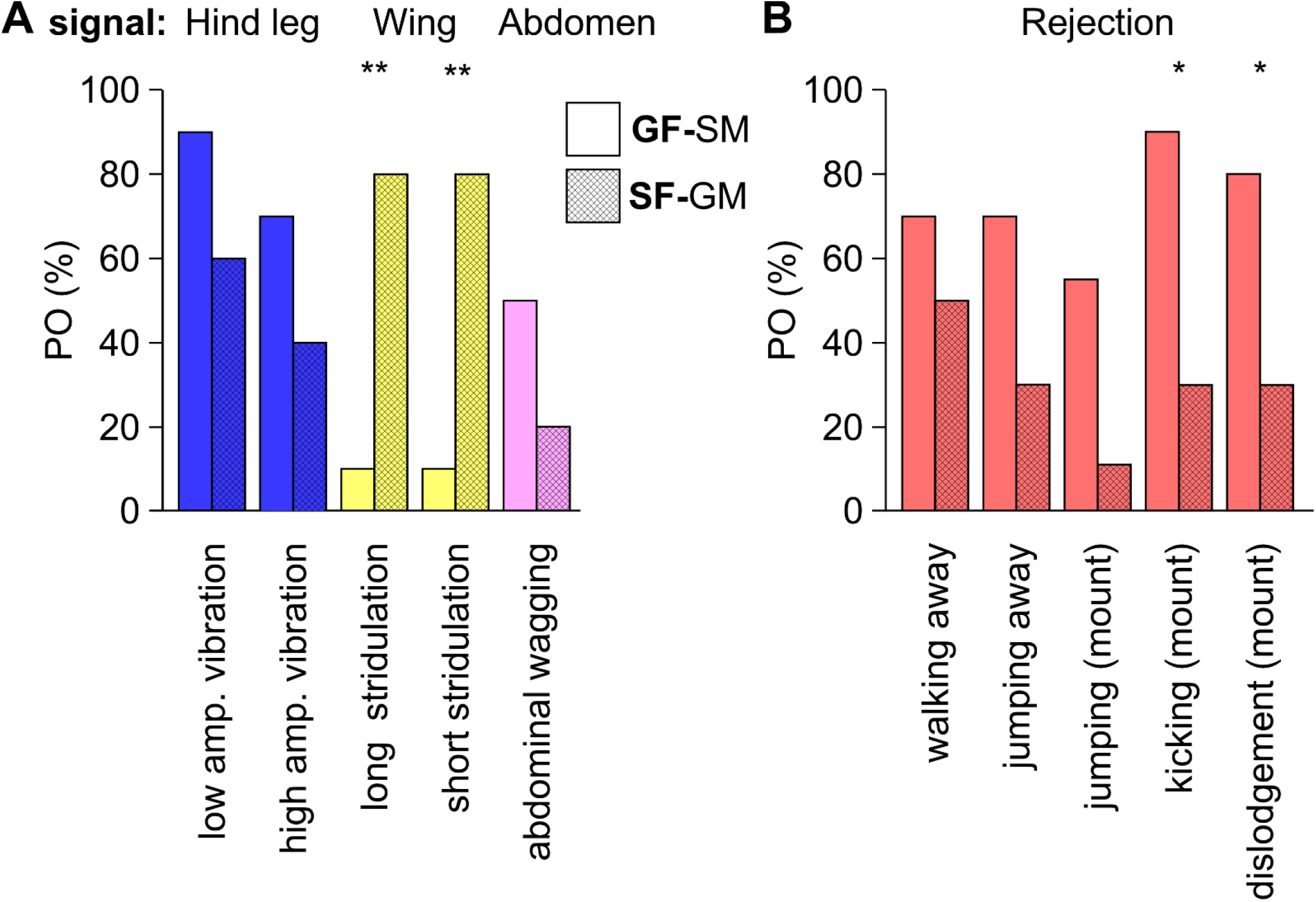
A. Pre-copulatory behavioral elements related to sexual behavior of the females in mixed-phase pair (GF-SM, Gregarious female and a solitarious male; or SF_GM, solitarious female and a gregarious male; n=10 for both). (mount) denotes after mounting, or during the mounting phase. Colors and bars as detailed in Fig 2. Raw data for the gregarious locusts are those found in Golov et al., 2018. Statistical significance: * p<0.05 ** p<0.01.

**Figure 6:**
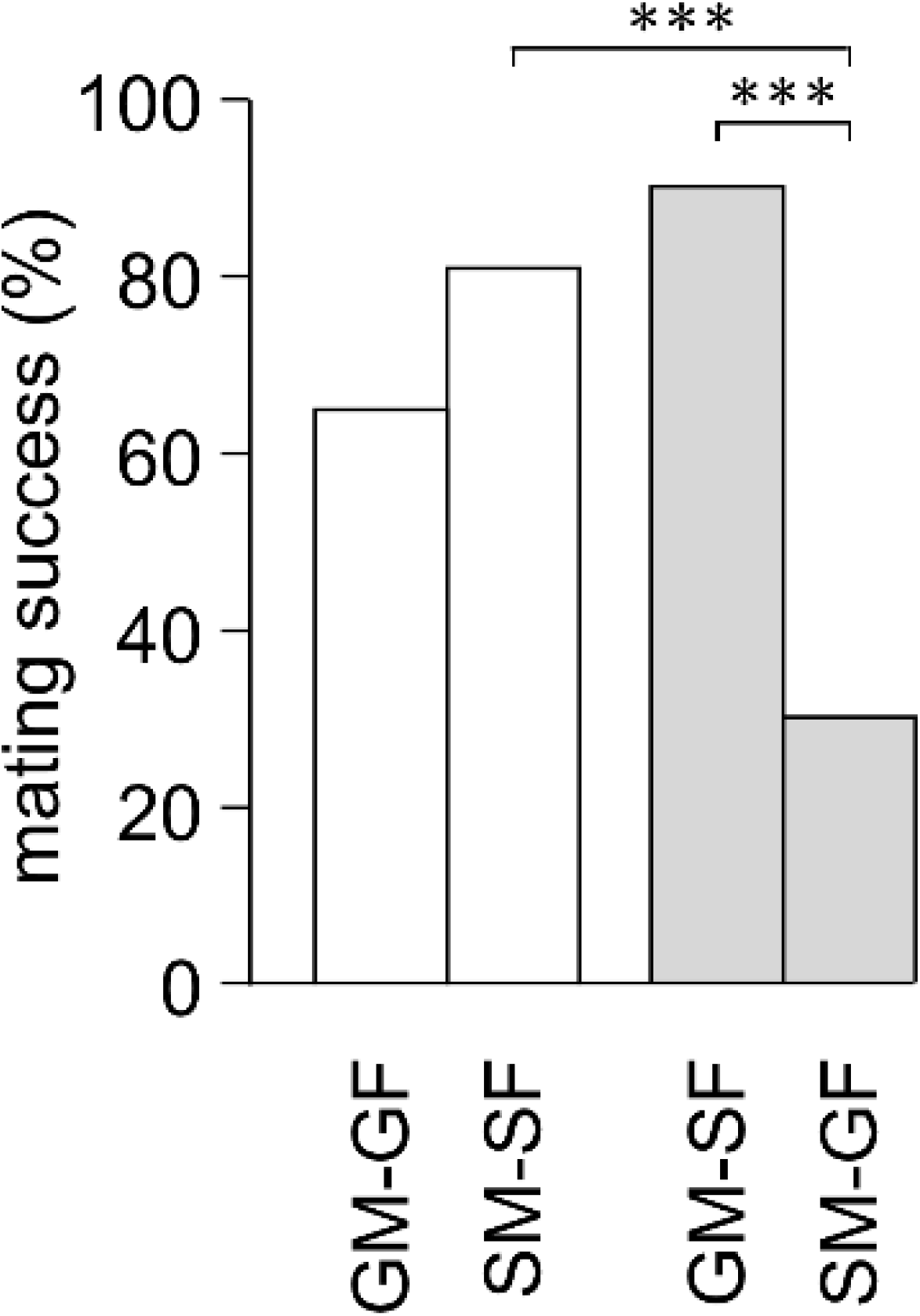
The probability of successful copulations in intra- and interphase pairs. GM and SM indicate gregarious and solitarious males, respectively; GF and SF indicate gregarious and solitarious females, respectively. Data for gregarious-gregarious couples were computed from the raw data found in Golov et al., 2018. Shaded bars denote data for the mixed/interphase pairs. *** p<0.001

## Discussion

Unique, species-specific, sexual behavior (i.e. courtship and mating-related behavioral patterns) is one of the hallmarks of a species, functioning both to defining and maintain species differentiation (Rundle & Nosil, 2005; Nosil, 2012). Hence, it is expected that courtship and mating-related behavioral patterns will be robust or highly consistent within the sexes of a species. However, ample research has revealed various sources of diversity or variation in sexual behavior and reproductive strategies within a species and, moreover, within sexes (Gross, 1996; Rhen & Crews, 2002).

Locust density-dependent phase polyphenism offers an extreme example of environmentally-induced plasticity, including very pronounced phase-dependent differences in behavior. The very different social environment of locusts in the gregarious or the solitarious phase is expected to result in pronounced differences in courtship and mating-related behavioral patterns. In a dense swarm of sexually mature gregarious locusts there is fierce competition among males. Field studies of *S. gregaria* have indicated that males outnumber females at the pairing and oviposition sites, and that there are severe struggles between males for the possession of females (Ellis & Ashall, 1957; Popov, 1958). This results in males displaying a highly active, dominant role in inter-sexual interactions (Golov *et al.*, 2018), with very basic or limited courtship, as also shown for the migratory locust (Oberlin, 1973), and which is assumed to be a characteristic of most Catantopinae (Otte, 1970). Strong et al. (1977) also suggested that limited male courtship is a characteristic of gregarious populations that experience strong intra-male competition. The gregarious females’ pre-copulatory behavior is mainly expressed by behavioral patterns aimed at rejecting the males (e.g. jumping and kicking, bending the abdomen: (Popov, 1958; Loher, 1959; Strong & Amerasinghe, 1977; Golov *et al.*, 2018), as known for most Catantopinae (Otte, 1970; Golov *et al.*, 2018).

In marked contrast, the solitarious locusts’ life history involves very different and major challenges in the path to successful mating and reproduction. These include: (1) locating a mate: while both male and female solitarious locusts may participate in mate location, it is accepted that it is mainly the males that are faced with the challenge of searching for and locating the females in a scarce population (Ely *et al.*, 2006; Maeno *et al.*, 2016); (2) sex recognition: this is not straightforward, since sexual dimorphism is very limited in the solitarious locusts compared to the gregarious phase (Norris, 1954; Pener, 1967; Pener & Simpson, 2009). Moreover, the overall behavior of the solitarious phase is characterized as cryptic – not supporting sex-related advertising; and (3) last but not least, there is a need to overcome the inherent tendency of solitarious locusts to repulse and avoid contact with conspecifics. These challenges should have led to: (a) the development of mechanisms dedicated to finding a mate. This important, interesting, and very little studied issue (e.g. Ely *et al.*, 2006) is, however, beyond the scope of the current work; (b) solitarious locusts should employ developed courtship behavior and inter-sex communication to serve in the process of sex recognition; and (c) the mating process of the solitarious pair should be enhanced by reinforcment signals serving to overcome their inherent repulsion tendency.

Indeed, our current findings well demonstrate the latter two points: Solitarious males were found to engage significantly more in stridulation behaviors (short and long), as well as in hind leg vibration, compared to gregarious males. Both behaviors may have an important role as signaling stimuli in inter-sex communication. Short and long stridulations were also demonstrated significantly more often by the solitarious compared to the gregarious females. Interestingly, these patterns of male courtship prior to mounting are not common among Catantopinae (Otte, 1970). Consistent with our hypothesis of a need for continuous reinforcement of the contact between the male and female, high level stridulation and leg vibration in the solitarious pairs were also extended into the mounting stage.

Evolution and sexual selection must have conveyed reproduction-related “benefits” for locusts in the solitarious phase that balance the above-detailed obstacles. These include greatly reduced intra-sex competition, and, as evident from our practically null observations of female rejection, little mating-related inter-sex conflict. It is possible, however, that the balance of obstacles vs. benefits differs for the solitarious desert locust males and females. The limited male-male competition probably leads the solitarious males to demonstrate limited intra-sex signaling, reduced assertiveness, and a general “low key” pre-copulatory behavior. On the other hand, solitarious females are tolerant, or even highly receptive during pre-mounting courtship, demonstrating very limited rejection-related behaviors throughout. Therefore, in respect to mating and reproduction, solitarious males and females may have different strategies. Such a sex-dependent strategy will have profound effects on the phase-related population dynamics, a point that should be further investigated both experimentally and in theoretical modeling.

The current research introduces another very important issue that should be considered in theoretical studies of locust population dynamics, and specifically phase-change dynamics (rarely addressed in theoretical modeling; Topaz *et al.*, 2012; Ariel & Ayali, 2015). Fundamental differences in pre-copulatory and courtship behavior are known to contribute to species-level reproductive isolation (e.g. Panhuis *et al.*, 2001; Masta & Maddison, 2002; Gray, 2005). Pre-mating isolation can result in significant levels of reproductive isolation, even in cases in which other reproductive isolation mechanisms are lacking (Nosil, 2012). The results of our experiments with the mixed, interphase pairs, clearly indicate that locusts of both phases tend to retain the basic features of their phase-specific pre-copulatory behavior when encountering a mate of the opposite phase. This point is further reinforced when remembering that we have specifically focused on behavioral patterns assumed to play a role in signaling and communication. Hence, has evolution and sexual selection result in limiting gene flow between sympatric solitarious and gregarious populations? A closer look at our results, again, suggests an asymmetry: i.e. that this may only be relevant to mixed pairs comprising a solitarious male and a gregarious female. Such a pair comprises “a male from Mars and a female from Venus” (in paraphrase of Gray, 1992), i.e. a pair with very different and far from complementary courtship and pre-copulatory behavioral phenotypes, to the extent that these profound differences may indeed serve to restrict inter-phase mating (as evident by our finding regarding low copulation success rates in such pairs). In marked contrast, the reciprocal pair, i.e. a gregarious male and a solitarious female, present a ‘tight match’ of two assertive and receptive counterparts, promoting rapid and high mating rates. Inter-phase sexual interactions may, therefore, be a case of asymmetrical sexual isolation (Arthur & Dyer, 2015). One outcome of the above may be an enhanced recruitment of solitarious females to a newly-formed gregarious group or a swarm passing by in an area inhabited by a solitarious locust population. Under specific conditions this can even constitute a mechanism for increasing the size of the swarm and escalating a locust outbreak.

The above findings may also assist in resolving a longstanding puzzle in the locust literature. Small populations of solitarious locusts should be at risk due to in-breeding and little overall gene flow. However, recent reports, investigating solitarious desert locust populations within a very large geographical range, have indicated a relatively high within-population genetic diversity and low between-population genetic structure (Chapuis *et al.*, 2014; Lazar *et al.*, 2016). In contrast, significant genetic differentiation among desert locust recession populations was also reported (Ibrahim *et al.*, 2000), suggesting that meta-population dynamics could produce genetic divergence among highly scattered populations (Ibrahim, 2001). In addition, in the migratory locust, *Locusta migratoria,* an homogenizing effect of past swarm events on solitarious populations was reported (Chapuis *et al.*, 2009). Our currently reported results, suggesting an additional layer of complexity in phase dynamics related to phase differences in mating and reproduction behavior as well as to intra-phase sex-dependent differential strategies, may aid in resolving such inconsistencies in solitarious desert locust population dynamics. For example, an inclination of solitary females to mate with gregarious males will allow a traveling group of gregarious locusts to serve as an instrument of genetic homogenization of remote solitarious populations. These ideas call for further research in the lab as well as in the field.

## Acknowledgments

We are grateful to Moshe Guershon for his assistance in the experiments and analysis and for live discussions. This work was funded by a grant from the Israel Ministry of Agriculture and Rural Development (891-0277-13). The work in its final stage was partially supported by the German Research Council (DFG; Grant RI 2728/2-1).

---

**Supplementary figure 1:**
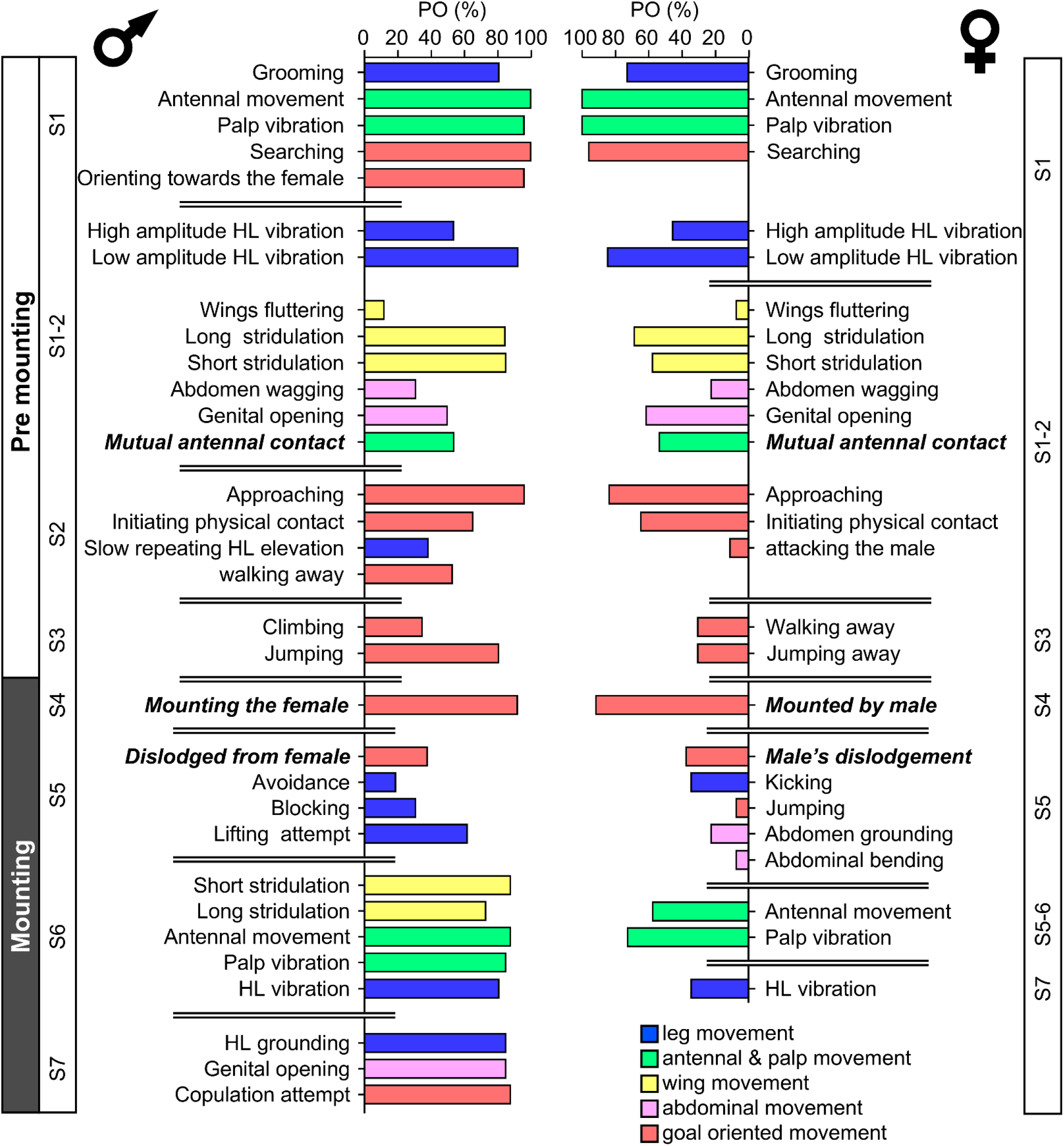
The precopulatory behavioral repertoire of the male (left) and female (right) desert locust in the solitary phase (Mean PO: The probability of an element to occur). The premounting and mounting behavioral elements are listed from step 1 to 7 (S1-S7) and color coded according to relevant body part. Behavioral elements that are shared and mutually exhibited by both sexes are presented in italic bold font.

**Supplementary figure 2:**
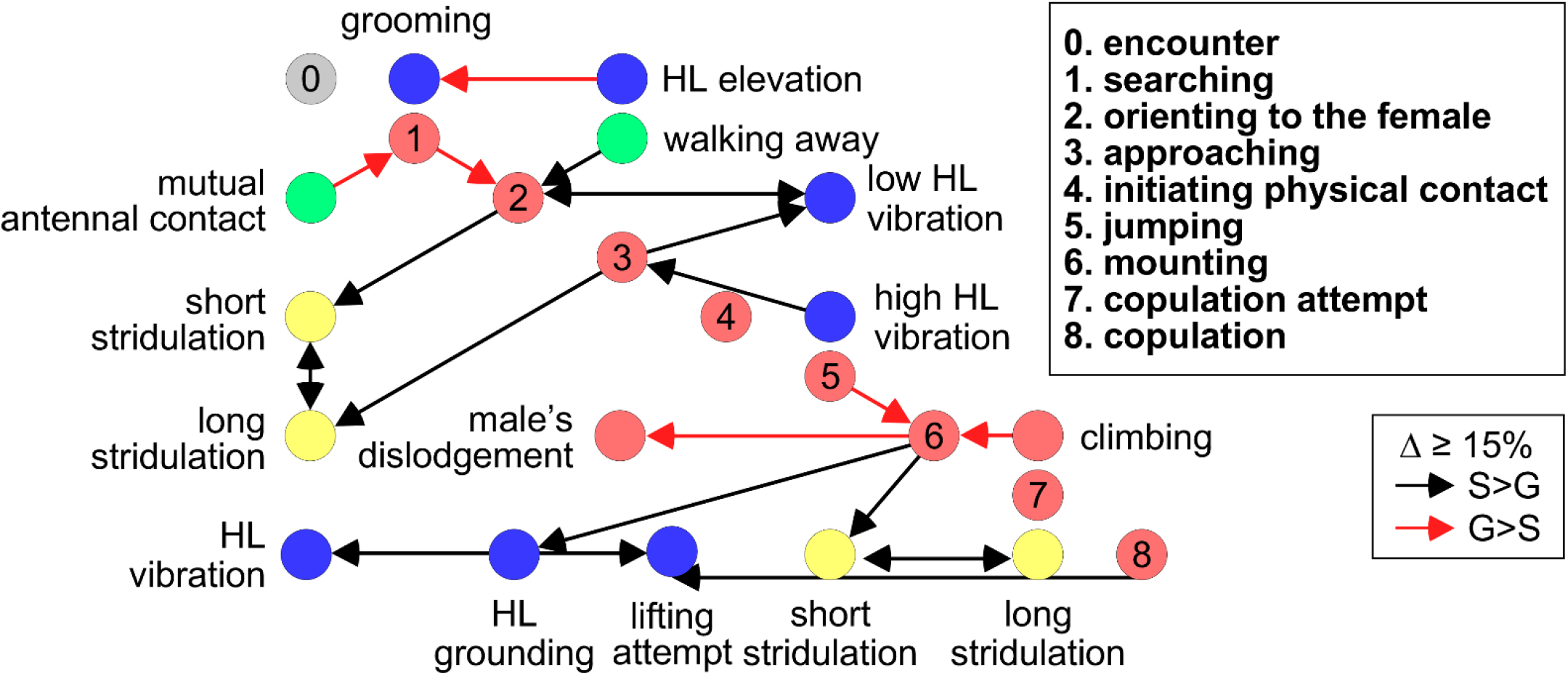
A comparison of the kinematics of the pre-copulatory behavior of solitary and gregarious male locusts; based on a matrix constructed for all transition probabilities (TP) between behavioral elements in each phase (as detailed in the Methods section and following Golov *et al.*, 2018). The color of the circles representing the different behavioral elements corresponds to the color index used in Fig. S1. Arrows represent the direction of a phase-dependent difference of 15% or more in the TP value between two elements, (red- higher in solitary, black-higher in gregarious locusts).

**Supplementary figure 3:**
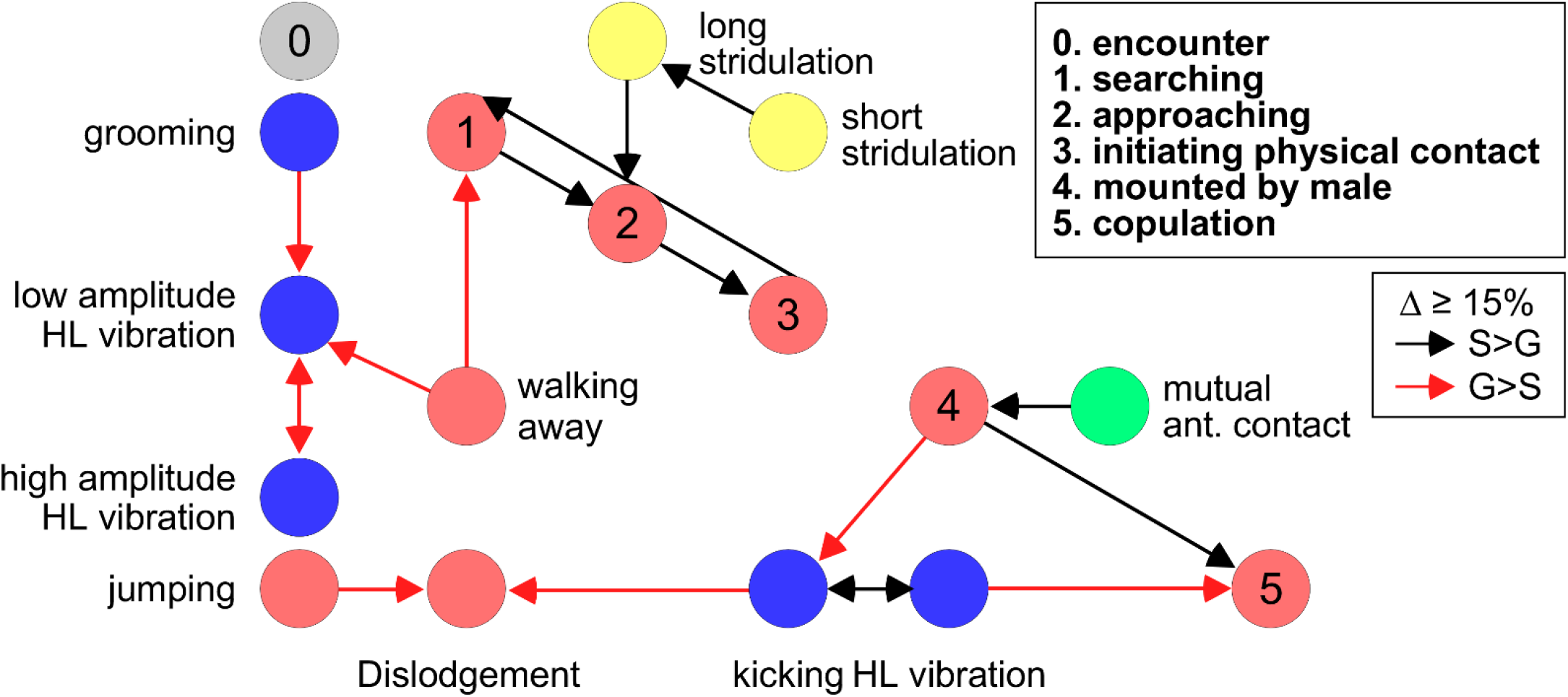
A comparison of the kinematics of the pre-copulatory behavior of solitary and gregarious female locusts; details as in Fig S2.

## Bibliography

Amerasinghe, F.P. (1978) Effects of J.H.I and J.H.III on yellowing, sexual activity and pheromone production in allatectomized male Schistocerca gregaria. Journal of Insect Physiology, 24, 603–611.

Andersson, M. & Iwasa, Y. (1996) Sexual selection. Trends in Ecology & Evolution, 11, 53–58.

Ariel, G. & Ayali, A. (2015) Locust collective motion and its modeling. PLoS Computational Biology, 11, e1004522.

Arthur, N.J. & Dyer, K.A. (2015) Asymmetrical sexual isolation but no postmating isolation between the closely related species Drosophila suboccidentalis and Drosophila occidentalis. BMC Evolutionary Biology, 15, 38.

Chapuis, M.-P., Loiseau, A., Michalakis, Y., Lecoq, M., Franc, A. & Estoup, A. (2009) Outbreaks, gene flow and effective population size in the migratory locust, Locusta migratoria: a regional-scale comparative survey. Molecular Ecology, 18, 792–800.

Chapuis, M.-P., Plantamp, C., Blondin, L., Pagès, C., Vassal, J.-M. & Lecoq, M. (2014) Demographic processes shaping genetic variation of the solitarious phase of the desert locust. Molecular Ecology, 23, 1749–1763.

Cullen, D.A., Cease, A.J., Latchininsky, A.V., Ayali, A., Berry, K., Buhl, J., et al. (2017) From molecules to management: mechanisms and consequences of locust phase polyphenism. In Advances in Insect Physiology. Academic Press, pp. 167–285.

Ellis, P.E. (1959) Learning and social aggregation in locust hoppers. Animal Behaviour, 7, 91–IN4.

Ellis, P.E. (1963) Changes in the social aggregation of locust hoppers with changes in rearing conditions. Animal Behaviour, 11, 152–160.

Ellis, P.E. & Ashall, C. (1957) Field Studies on diurnal Behaviour, Movement and Aggregation in the Desert Locust (Schistocerca gregaria Forskǻl). Field Studies on diurnal Behaviour, Movement and Aggregation in the Desert Locust (Schistocerca gregaria Forskǻl).

Ely, S.O., Mahamat, H., Njagi, P.G.N., Bashir, M.O., El-Amin, S.E.-T. & Hassanali, A. (2006) Mate location mechanism and phase-related mate preferences in solitarius desert locust, Schistocerca gregaria. Journal of Chemical Ecology, 32, 1057–1069.

Farrow, R.A. (1979) Population dynamics of the australian plague locust, chortoicetes terminifera (walker), in central western new south wales. I. reproduction and migration in relation to weather. Australian journal of zoology, 27, 717.

Friard, O. & Gamba, M. (2016) BORIS: a free, versatile open-source event-logging software for video/audio coding and live observations. Methods in ecology and evolution / British Ecological Society, 7, 1325–1330.

Geva, N., Guershon, M., Orlova, M. & Ayali, A. (2010) Memoirs of a locust: density-dependent behavioral change as a model for learning and memory. Neurobiology of Learning and Memory, 93, 175–182.

Golov, Y., Rillich, J., Harari, A. & Ayali, A. (2018) Precopulatory behavior and sexual conflict in the desert locust. PeerJ, 6, e4356.

Gray, D.A. (2005) Does courtship behavior contribute to species-level reproductive isolation in field crickets? Behavioral Ecology, 16, 201–206.

Gray, J. (1992) Men are from Mars, women are from Venus: A practical guide for improving communication and getting what you want in your relationships. 1st ed. HarperCollins, New York, NY.

Gross, M.R. (1996) Alternative reproductive strategies and tactics: diversity within sexes. Trends in Ecology & Evolution, 11, 92–98.

Ibrahim, K.M. (2001) Plague dynamics and population genetics of the desert locust: can turnover during recession maintain population genetic structure? Molecular Ecology, 10, 581–591.

Ibrahim, K.M., Sourrouille, P. & Hewitt, G.M. (2000) Are recession populations of the desert locust (Schistocerca gregaria) remnants of past swarms? Molecular Ecology, 9, 783–791.

Inayatullah, C., El Bashir, S. & Hassanali, A. (1994) Sexual Behavior and Communication in the Desert Locust, Schistocerca gregaria (Orthoptera: Acrididae): Sex Pheromone in Solitaria. Environmental Entomology, 23, 1544–1551.

Injeyan, H.S. & Tobe, S.S. (1981) Phase polymorphism in Schistocerca gregaria: Reproductive parameters. Journal of Insect Physiology, 27, 97–102.

Lazar, M., Piou, C., Doumandji-Mitiche, B. & Lecoq, M. (2016) Importance of solitarious desert locust population dynamics: lessons from historical survey data in Algeria. Entomologia experimentalis et applicata, 161, 168–180.

Loher, W. (1959) Contributions to the study of the sexual behaviour of schistocerca gregaria forskål (orthoptera: acrididae). Proceedings of the Royal Entomological Society of London. Series A, General Entomology, 34, 49–56.

Loher, W. (1961) The chemical acceleration of the maturation process and its hormonal control in the male of the desert locust. Proceedings of the Royal Society B: Biological Sciences, 153, 380–397.

Maeno, K.O., Ould Ely, S., Nakamura, S., Abdellaoui, K., Cissé, S., Jaavar, M.E.H., et al. (2016) Daily microhabitat shifting of solitarious-phase Desert locust adults: implications for meaningful population monitoring. SpringerPlus, 5, 107.

Mahamat, H., Hassanali, A., Odongo, H., Torto, B. & El-Bashir, E.-S. (1993) Studies on the maturation-accelerating pheromone of the desert locustSchistocerca gregaria (Orthoptera: Acrididae). Chemoecology, 4, 159–164.

Masta, S.E. & Maddison, W.P. (2002) Sexual selection driving diversification in jumping spiders. Proceedings of the National Academy of Sciences of the United States of America, 99, 4442–4447.

Nishide, Y. & Tanaka, S. (2012) Yellowing, morphology and behaviour in sexually mature gynandromorphs of the desert locust*Schistocerca gregaria*. Physiological entomology, 37, 379–383.

Norris, M.J. (1954) Sexual maturation in the desert locust (Schistocerca gregariaForskål) with special reference to the effects of grouping. Anti-Locust Bulletin, 18, 1–44.

Norris, M.J. (1962) Group effects on the activity and behaviour of adult males of the desert locust (Schistocerca gregaria Forsk.) in relation to sexual maturation. Animal Behaviour, 10, 275–291.

Nosil, P. (2012) Ecological Speciation. OUP Oxford, Oxford.

Nosil, P., Vines, T.H. & Funk, D.J. (2005) Reproductive isolation caused by natural selection against immigrants from divergent habitats. Evolution, 59, 705–719.

Obeng-Ofori, D., Torto, B. & Hassanali, A. (1993) Evidence for mediation of two releaser pheromones in the aggregation behavior of the gregarious desert locust, Schistocerca gregaria (forskal) (Orthoptera: Acrididae). Journal of Chemical Ecology, 19, 1665–1676.

Oberlin, U.P. (1973) Verhaltensbiologische Studien an der europaeischen Wanderheuschrecke Locusta Migratoria L. Entomol Gesell Basel, 23, 12–23.

Otte, D. (1970) A comparative study of communicative behaviour in grasshoppers. Misc Publ Mus Zool Univ Mich, 141, 1–168.

Panhuis, T.M., Butlin, R., Zuk, M. & Tregenza, T. (2001) Sexual selection and speciation. Trends in Ecology & Evolution, 16, 364–371.

Pener, M.P. (1965) On the influence of corpora allata on maturation and sexual behaviour of Schistocerca gregaria. Journal of zoology, 147, 119–136.

Pener, M.P. (1967) Effects of allatectomy and sectioning of the nerves of the corpora allata on oöcyte growth, male sexual behaviour, and colour change in adults of Schistocerca gregaria. Journal of Insect Physiology, 13, 665–684.

Pener, M.P. (1976) The differential effect of the corpora allata on male sexual behaviour in crowded and isolated {ILocusta migratoria migratorioides} (R & F.) males. Acrida, 5, 189–206.

Pener, M.P. & Simpson, S.J. (2009) Locust phase polyphenism: an update. In Advances in insect physiology volume 36, Advances in insect physiology. Elsevier, pp. 1–272.

Popov, G.B. (1958) Ecological Studies on Oviposition by Swarms of the Desert Locust (Schistocerca gregaria Forskal) in eastern Africa. Ecological Studies on Oviposition by Swarms of the Desert Locust (Schistocerca gregaria Forskal) in eastern Africa., 1–70.

Ramirez-Romero, R., Garibay-Benítez, D., Vargas-Ponce, O., Joyce, A. & Bernal, J.S. (2017) Do assortative mating and immigrant inviability help maintain population genetic structuring of an herbivore on a crop and a wild relative? Insect science (Online).

Rhen, T. & Crews, D. (2002) Variation in reproductive behaviour within a sex: neural systems and endocrine activation. Journal of Neuroendocrinology, 14, 517–531.

Roessingh, P., Simpson, S.J. & James, S. (1993) Analysis of Phase-Related Changes in Behaviour of Desert Locust Nymphs. Proceedings of the Royal Society B: Biological Sciences, 252, 43–49.

Rundle, H.D. & Nosil, P. (2005) Ecological speciation. Ecology Letters, 8, 336–352.

Strong, L. & Amerasinghe, F.P. (1977) Allatectomy and sexual receptivity in females of Schistocerca gregaria. Journal of Insect Physiology, 23, 131–135.

Tanaka, S., Maeno, K., Ould Mohamed, S., Ould Ely, S. & Babah Ebbe, M.A. (2010) Upsurges of desert locust populations in Mauritania: Body coloration, behavior and morphological characteristics. Applied entomology and zoology, 45, 641–652.

Tanaka, S. & Zhu, D.-H. (2003) Phase-Related Differences in Mating Strategy of a Locust (Orthoptera: Acrididae). Acta Neophilologica, 96, 498–502.

Topaz, C.M., D’Orsogna, M.R., Edelstein-Keshet, L. & Bernoff, A.J. (2012) Locust dynamics: behavioral phase change and swarming. PLoS Computational Biology, 8, e1002642.

Uvarov, B. (1977) Grasshoppers and locusts. A handbook of general acridology Vol. 2. Behaviour, ecology, biogeography, population dynamics. cabdirect.org.

Uvarov, B.P. (1966) Phase polymorphism. In Grasshoppers and locusts. Cambridge university press, Cambridge, pp. 332–386.

Whitman, D.W. (1990) Grasshopper chemical communication. In Biology of Grasshoppers (ed. Chapman, R.F. & Joern, A.). Wiley, New York, pp. 357–391.

Wybrandt, G.B. & Andersen, S.O. (2001) Purification and sequence determination of a yellow protein from sexually mature males of the desert locust, Schistocerca gregaria. Insect Biochemistry and Molecular Biology, 31, 1183–1189.

